# A Deep Learning Framework for Biomarker Segmentation and Classification in Traumatic Brain Injury

**DOI:** 10.64898/2026.07.09.737265

**Authors:** Rohan Dash, Karthick Mayilsamy, Ryan Green, Yu Sun, Shyam S Mohapatra, Subhra Mohapatra

**Affiliations:** Department of Molecular Medicine, University of South Florida; Department of Internal Medicine, Morsani College of Medicine, University of South Florida; Bellini College of Medicine, University of South Florida; James A Haley Veterans Hospital, Tampa, FL

**Keywords:** machine learning, classification, segmentation, multimodal, traumatic brain injury, microglia, astrocytes

## Abstract

Traumatic brain injury (TBI) triggers widespread biomarker activation, including astrocytic markers such as glial fibrillary acidic protein (GFAP) and microglia markers such as ionized calcium-binding adapter molecule 1 (IBA1). Quantifying and analyzing these biomarkers are critical for understanding injury impact; however, current methods are labor-intensive and time-consuming. In this study, we propose an automated deep learning framework for dual-biomarker segmentation and TBI classification using GFAP and IBA1 immunofluorescent images. Four U-Net variants: Baseline U-Net, U-Net++, MANet, and LinkNet were trained for segmentation. Three classification models, ResNet50, Swin_T, and MaxViT, were trained to distinguish TBI from control images under single- and dual-biomarker conditions. The baseline U-Net achieved the highest segmentation Dice score for GFAP (0.9259), while the U-Net++ achieved the highest Dice score for IBA1 (0.9676). Trained segmentation models demonstrated significantly better performance compared to QuPath alternatives. While GFAP alone supported high classification accuracy, IBA1 alone was less effective. Multimodal fusion of GFAP and IBA1 significantly improved classification performance across all models, with Swin_T achieving the highest overall accuracy (0.9489), and ResNet50 achieving the highest F1-score (0.9499). These findings demonstrate that integrating complementary biomarkers enhances automated TBI classification, and deep learning offers a robust alternative to manual analysis for immunofluorescent brain injury imaging. This framework is scalable to additional biomarkers and injury models, offering a reproducible approach to accelerate biomarker research.

## 1. Background

Traumatic brain injury (TBI) is a leading cause of neurological disability worldwide, affecting over 50 million individuals annually [1]. TBI can lead to immediate impairments such as loss of consciousness, cerebral hypoxia [2], headaches, seizures, and behavioral abnormalities [3], and in severe cases, mortality [4]. Long-term consequences often include persistent motor dysfunction and cognitive deficits [5]. Beyond individual patient suffering, especially in high-risk populations [6], [7], TBI imposes a significant economic burden, with global costs exceeding $400 billion annually [8]. Given its clinical, societal and economic impact, understanding TBI at the cellular level is critical.

Following TBI, glial cells in the central nervous system undergo rapid activation, characterized by distinct morphological and molecular changes. This activation is associated with increased expression of specific biomarkers. Evidence suggests that microglial and astrocyte activation influences the brain’s response to and recovery after TBI [9]. Ionized calcium-binding adapter molecule 1 (IBA1) and glial fibrillary acidic protein (GFAP) are widely used immunohistochemical markers to microglia and astrocytes, respectively. These biomarkers provide invaluable insight on neuroinflammatory responses to injury. GFAP remains one of the most frequently analyzed biomarkers in neuroscience [10], whereas IBA1 is a reliable marker that shows robust expansion following TBI [11]. Because these biomarkers are visualized through histological imaging, quantitative image analysis is key to understanding cellular responses to TBI.

Manual or semi-automated analysis of such images is labor-intensive and inefficient [12], underscoring the need for automated and reproducible computational methods. Deep learning has demonstrated strong performance in biomedical image classification and segmentation [13], [14]. In computer vision, convolutional neural networks (CNNs) dominate due to their ability to deliver high accuracy with relatively low computational cost [15], [16]. Originally developed for general image analysis, these models have been widely adapted for medical imaging [17], [18]. For segmentation tasks, U-Net architectures have become the standard in medical imaging because of their encoder-decoder design and consistently high performance [19], [20], [21].

Recent studies have applied these techniques to glial histology. For example, one study combined directional filters with CNNs to segment GFAP-labeled astrocytes in fluorescent images, outperforming traditional methods [22]. Similarly, another study developed an automated microglia detection method using object detection-based CNNs, achieving 94% precision, 91% recall, and 92% F1-score [23]. Deep CNNs have also been used for astrocyte segmentation, performing comparably to human experts with an 81% F1-score [24].

Despite these advances, most studies have analyzed astrocytes and microglia independently, potentially missing insights into the coordinated activation of these glial populations. Given their complementary roles in neuroinflammation, a multimodal approach may improve classification accuracy by combining GFAP and IBA1 within a unified deep learning framework. Multimodal fusion methods in biomedical imaging have been shown to outperform single channel models [25], supporting the hypothesis that incorporating multiple biomarkers yields more robust models.

In this study, we introduce a novel deep learning pipeline that (i) trains and compares multiple U-Net variants to segment immunofluorescent GFAP and IBA1 images, (ii) trains and compares different classifier architectures to predict TBI status, and (iii) compares unimodal (singular biomarker) versus multimodal (dual biomarker) approaches. We assess performance using K-fold cross-validation and standardized metrics. We hypothesize that a multimodal fusion of GFAP and IBA1 histological images via deep learning will improve TBI classification accuracy over unimodal models. To our knowledge, this is the first systematic evaluation of dual-biomarker segmentation and classification for TBI histology using deep learning. Our framework provides an automated, reproducible method for biomarker analysis, reducing manual effort and enabling extension to additional biomarkers and injury models, addressing a critical challenge in biomedical imaging.

## 2. Materials and Methods

The overall methodology is illustrated in Fig. 1. The process begins with acquiring images from mouse brain samples and preprocessing with Python and Fiji (ImageJ). Preprocessed images are then used in a two-stage machine learning pipeline: (i) image segmentation and (ii) image classification. A total of twelve segmentation-classification model combinations were trained and evaluated. Segmentation models are trained separately for IBA1 and GFAP, while classification models have three tasks: IBA1 only, GFAP only, and combined GFAP + IBA1.

**Fig. 1.**
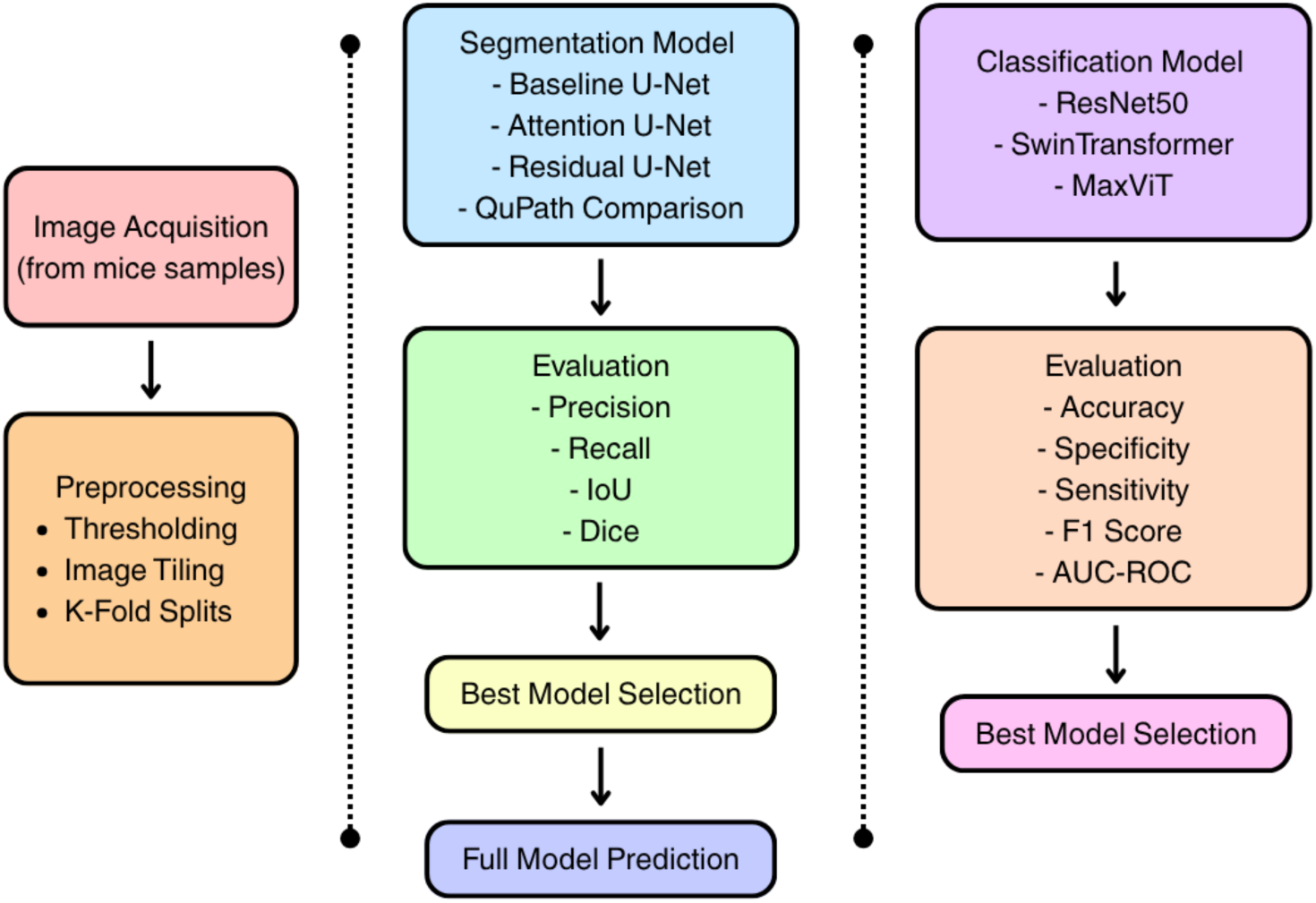
Overall approach to Image acquisition, preprocessing and model selection and evaluation.

### 2.1 Data Acquisition

All procedures were approved by the Institutional Animal Care and Use Committee (IACUC) of the University of South Florida and conducted in accordance with NIH guidelines for the Care and Use of Laboratory Animals. Male C57BL/6 mice (12–14 weeks old) were housed under a 12-h light/dark cycle with ad libitum access to food and water. Repeated closed-head TBI was induced following previously established protocols [26].

Mice were euthanized 7 days post-injury, deeply anesthetized, and transcardially perfused with PBS followed by 4% paraformaldehyde. Brains were post-fixed in 30% sucrose for 48–72h at 4 °C, frozen, and coronally sectioned at 30 µm using a cryostat. Sections were processed for immunofluorescence staining. Microglial activation was assessed using an anti-Iba1 antibody (rabbit anti-Iba1, Wako, 019-19741; 1:1000). Astrogliosis was evaluated using an anti-GFAP antibody (chicken anti-GFAP, Invitrogen, AB5541; 1:1000). Secondary antibodies included goat anti-rabbit Alexa Fluor 594 (Invitrogen, A11008; 1:1000) and goat anti-chicken Alexa Fluor 488 (Invitrogen, AB11039; 1:1000). Nuclei were counterstained with DAPI (1:10,000; Fisher Scientific). An average of 3–5 brain sections per animal was analyzed with two images captured per section, yielding 10 images per animal from the perilesional cortex surrounding the injury site. Immunoreactivity for Iba1 and GFAP was quantified using NIH ImageJ software. All images were acquired using identical exposure and digital gain settings to minimize variability. Images were collected as part of three groups: GFAP, IBA1, and paired GFAP-IBA1, each with control and TBI classes. All images are sized 1360 × 1024 pixels. Table 1 summarizes image counts, with paired counts reflecting the total number of images (49 control and 50 TBI pairs).

**Table 1.** Image counts.

| **Marker** | **Control** | **TBI** |
| --- | --- | --- |
| GFAP | 99 | 75 |
| IBA1 | 276 | 383 |
| Paired GFAP-IBA1 | 98 | 100 |

### 2.2 Data Preprocessing

Preprocessing was necessary to prepare raw immunofluorescence images for training deep learning models on segmentation and classification tasks [27]. All images were processed using Fiji (ImageJ). Each raw image was first converted to 8-bit grayscale and thresholded using Fiji’s Default automatic algorithm in black-and-white mode with a dark background. To standardize thresholding, ten representative images per biomarker (five Control and five TBI) were randomly selected and manually thresholded using the same algorithm. The resulting threshold values were then averaged and uniformly applied to all images within each biomarker group. After thresholding, all binary masks were manually verified and visually inspected by a trained researcher to ensure accurate correspondence between each mask and its immunofluorescent image.

Binary masks were generated with pixel values ranging from 0 to 255, then normalized and rescaled to a range of 0 to 1. No additional intensity normalization or background correction was applied. To train the models most effectively, K-fold cross validation, a widely utilized method of training machine learning models [28], [29] was utilized, where K = 5 for this methodology. K-fold cross validation is considered beneficial over traditional train-test fold splits to allow for a more robust estimate of its performance. The dataset was partitioned into five mutually exclusive folds, with all images originating from the same animal kept within the same fold to prevent data leakage. Across five training cycles, the folds were rotated so that, in each cycle, three folds were used for training, one fold was used for validation, and one fold was used for testing. This procedure was then repeated, allowing for ten separate trials. Following normalization and fold splits, images were tiled for suitable model input [30], [31] as seen in Figure 2.

**Figure 2.**
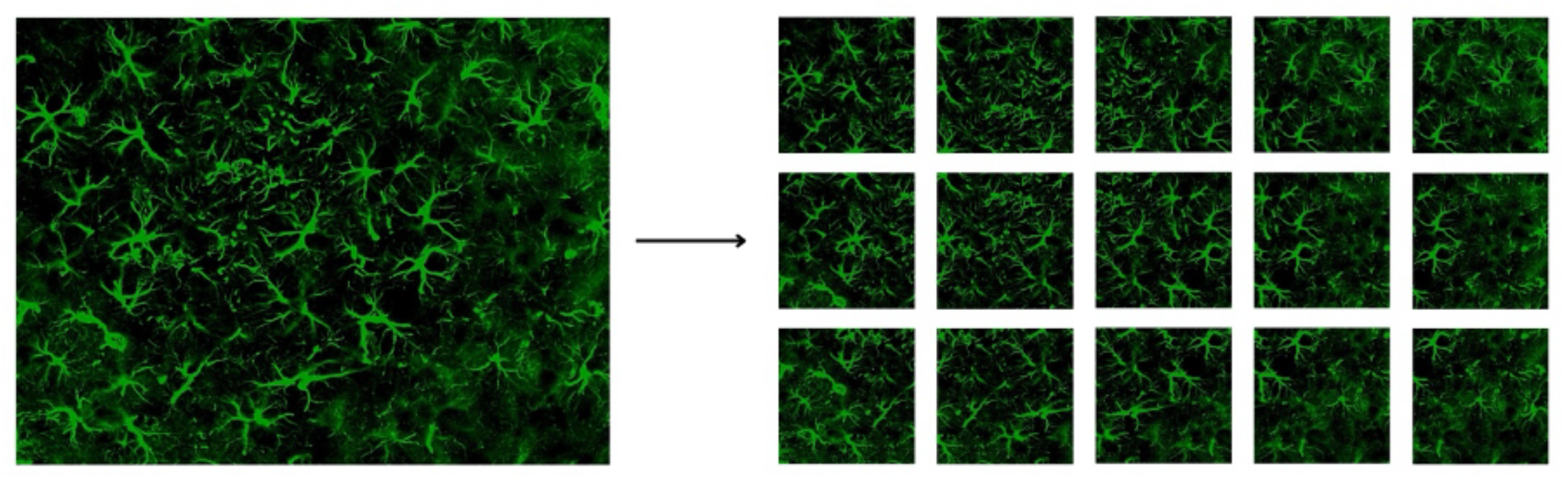
Tiling allowed for one 1360×1024 image to be converted to 15 512×512 images. Subsequently, segmentation models were trained, validated, and tested with tiled images, which were then stitched back for classification tasks.

Because the original dimensions for all images were 1360 × 1024 pixels, each image was tiled into overlapping 512 × 512 pixel patches [32] with a 50% overlap between adjacent tiles (256 pixel stride). This procedure produced 15 tiles per image, using the following horizontal ranges: 0–512 px, 256–768 px, 512–1024 px, 768–1280 px, and 848–1360 px for the final tile, which matched the remaining image width without any padding. Vertically, tiles spanned 0–512 px, 256–768 px, and 512–1024 px. By doing splits before tiling, we were able to ensure no data leakage between different folders.

### 2.3 U-Net Training Setup

Segmentation was performed independently for each biomarker (GFAP and IBA1), with separate models trained for each; no pairing or mixing was applied. Training, validation, and testing followed the same five-fold cross-validation structure from Section 2.2, ensuring all images from the same animal remained within the same fold to prevent leakage. Binary masks derived in Section 2.2 served as ground truth for training. Each model was trained for 50 epochs per fold using the Adam optimizer and a learning rate of 1×10^-4^. The training batch size was set to 64, and automatic mixed precision was enabled to improve stability and speed.

Four different U-Nets were utilized: the standard U-Net architecture [19], U-Net++, MANet, and LinkNet. U-Net++ is an improved version of the Standard U-Net by using convolution blocks first before moving features from the encoder to the decoder [33]. MANet uses multi-scale attention mechanisms in its encoder for contextual information [34]. LinkNet uses skip connections that connect the encoder and decoder [35]. ResNet18, which is known for its skip connections [36], [37] is used as the encoder for all four models.

To address class imbalance and improve segmentation stability, a combined BCE–Dice loss was used consisting of equal weights (0.5 each) for Binary Cross-Entropy with Logits and soft Dice loss, with a smoothing constant of 1×10^-6^. Equation 1 expresses this formally below.

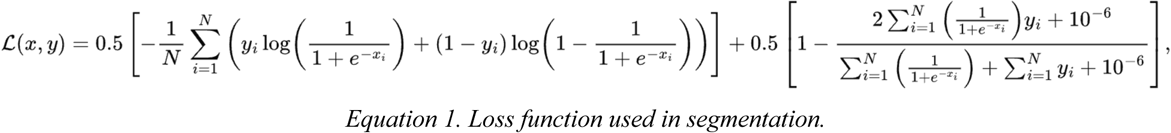

Data augmentation was applied only to training data and was limited to biologically plausible geometric transformations: horizontal flip, vertical flip, and a random rotation of 0°, 90°, 180°, or 270°. During training, validation Dice and validation loss were computed at each epoch. The checkpoint with the highest validation Dice score within the 50-epoch window was selected as the final model for that fold. Evaluation was then conducted on all ten folds.

To compare the effectiveness of using specific U-Net models, we trained three pixel-level classifiers within QuPath using the OpenCV machine learning library: Artificial Neural Network, Random Forest, and Logistic Regression. GeoJSON masks were created for positive biomarker regions, and a background annotation was created by subtracting biomarker regions from the full image. A five-fold setup was created twice to replicate the number of trials with the U-Nets. Because validation is not possible within QuPath machine learning models, we trained each model on four folds and tested on one fold. All default values were used, with a resolution downsample of 1.0 and default multiscale features in two dimensions. Model performance was evaluated by comparing the models’ predicted biomarker annotations with the annotated ground-truth masks. Predicted and reference annotations were converted to geometric regions, from which true-positive, false-positive, and false-negative areas were computed for metrics.

### 2.4 Classification CNN Training Setup

Following segmentation, a classification stage was implemented to train models to determine whether each image originated from a TBI or Control sample. This stage assessed whether multimodal information from GFAP and IBA1 improved performance over a single biomarker.

Recent advances in image classification highlight three strong model types: convolutional neural networks [38], vision transformers [39], [40] and hybrid models combining both [41]. Selected models and their parameters are listed in Table 2. Models were chosen with similar parameter counts, and preprocessing differences (e.g., image size) were considered during training. All models were instantiated using the timm library with ImageNet-pretrained weights, modified to accept either three or six input channels depending on the experiment. The output layer was set to two classes: Control and TBI. For the classification task, we utilized the immunofluorescent images as the input for chosen models. This approach is consistent with established biological analysis pipelines, where classification is performed on raw fluorescent channel images as key features for classification are encoded in the spatial features of the biomarkers [42], [43]. This also avoids classic shortcut learning issues with binary image masks.

**Table 2.** Model size and type. All three models had a slightly greater number of parameters (from when modified to accept six input channels).

| Model Name | Parameters (3-channel) | Parameters (6-channel) | Model Type |
| --- | --- | --- | --- |
| ResNet50 | 23,512,130 | 23,521,538 | CNN |
| Swin_T | 27,520,892 | 27,525,500 | ViT |
| MaxViT | 30,404,554 | 30,406,282 | Hybrid |

All classification models were trained using the same two-time five-fold cross-validation framework as segmentation, ensuring consistency in fold assignments and preventing data leakage. Pre-trained models were initialized with ImageNet weights and fine-tuned for respective biomarker classification tasks. No backbone layers were frozen; all weights were allowed to update during training. For multimodal experiments, paired GFAP-IBA1 images remained in the same fold to maintain biological correspondence. The same biologically relevant augmentations described in Section 2.2 were applied again at this stage.

Models were trained for 60 epochs in each fold. The AdamW optimizer is utilized with a learning rate of 1×10^-4^ and weight decay of 1×10^-4^. A patience of 15 epochs was utilized to end training early if convergence had already been achieved. Images were resized to 224 x 224 pixels. Equation 2 demonstrates the cross-entropy loss function that was used. The batch size was set to 32, based on GPU memory constraints. During training, validation loss and accuracy were monitored at each epoch. All model weights and training logs were saved to ensure full reproducibility. This classification framework allowed direct evaluation of unimodal versus multimodal learning. Evaluation across each fold was done twice using two different random seeds.

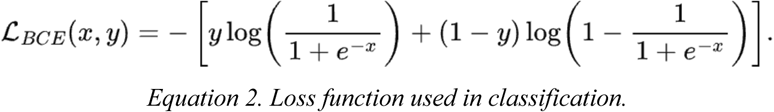

### 2.5 Statistics and Data Analysis

Model performance was evaluated using standard computer vision metrics for segmentation and classification. For segmentation, four primary metrics were used to quantify performance and the difference between the predicted masks and the ground-truth masks: Dice coefficient, Intersection over Union (IoU), precision, and recall. Probability maps were thresholded at 0.5 to obtain binary masks. All segmentation metrics were computed tile-wise on the held-out test fold of each split. Segmentation checkpoints were selected based on the highest validation Dice score during training. For classification, model outputs were predicted as either Control or TBI using a 0.5 decision threshold for the TBI class. Classification performance was evaluated with accuracy, sensitivity (true positive rate), specificity (true negative rate), F1-score, and AUC-ROC (area under the receiver operating characteristic curve). All classification metrics were computed on the held-out test fold of each split and reported as the mean ± standard deviation, calculated across the ten test-fold results (five folds, repeated twice).

## 3. Results

### 3.1 GFAP Segmentation

The GFAP segmentation results in Table 3 demonstrate strong performance across all four architectures, with U-Net obtaining the highest Dice coefficient of 0.9259 ± 0.0229 and an IoU of 0.8670 ± 0.0389, outperforming all three other U-Nets. Precision and recall values are consistently high across all models, indicating reliable identification of positive pixels, with the highest of 0.9291 from U-Net++. Notably, while LinkNet exhibits lower Dice and IoU scores, its recall is better than that of MANet, indicating strong performance with positive regions despite relatively inconsistent boundary prediction. LinkNet demonstrated the lowest IoU among the four models. Representative segmentation examples for both classes are shown in Fig. 3.

**Figure 3.**
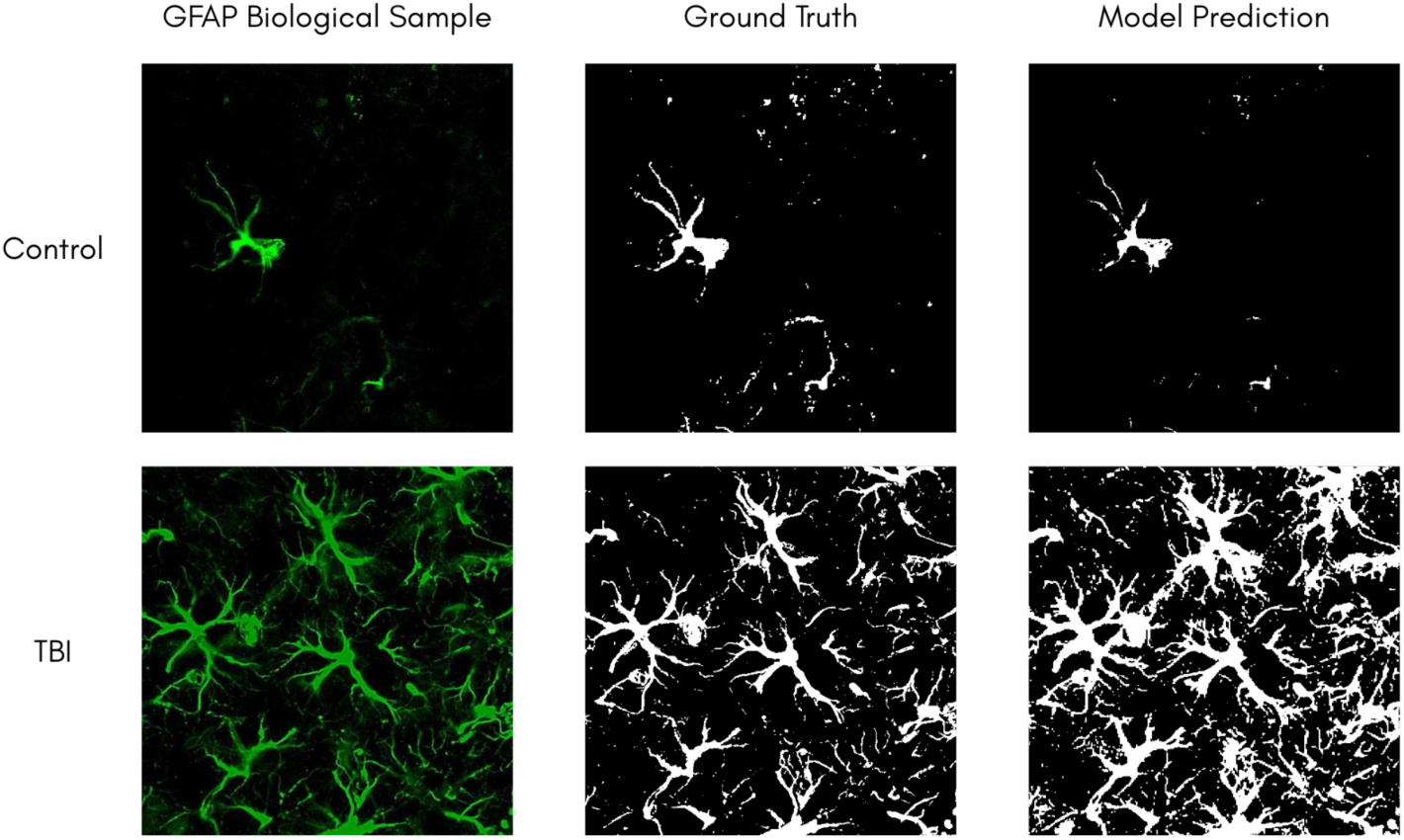
GFAP segmentation prediction for TBI and Control, shown alongside original tile and ground truth.

**Table 3.**
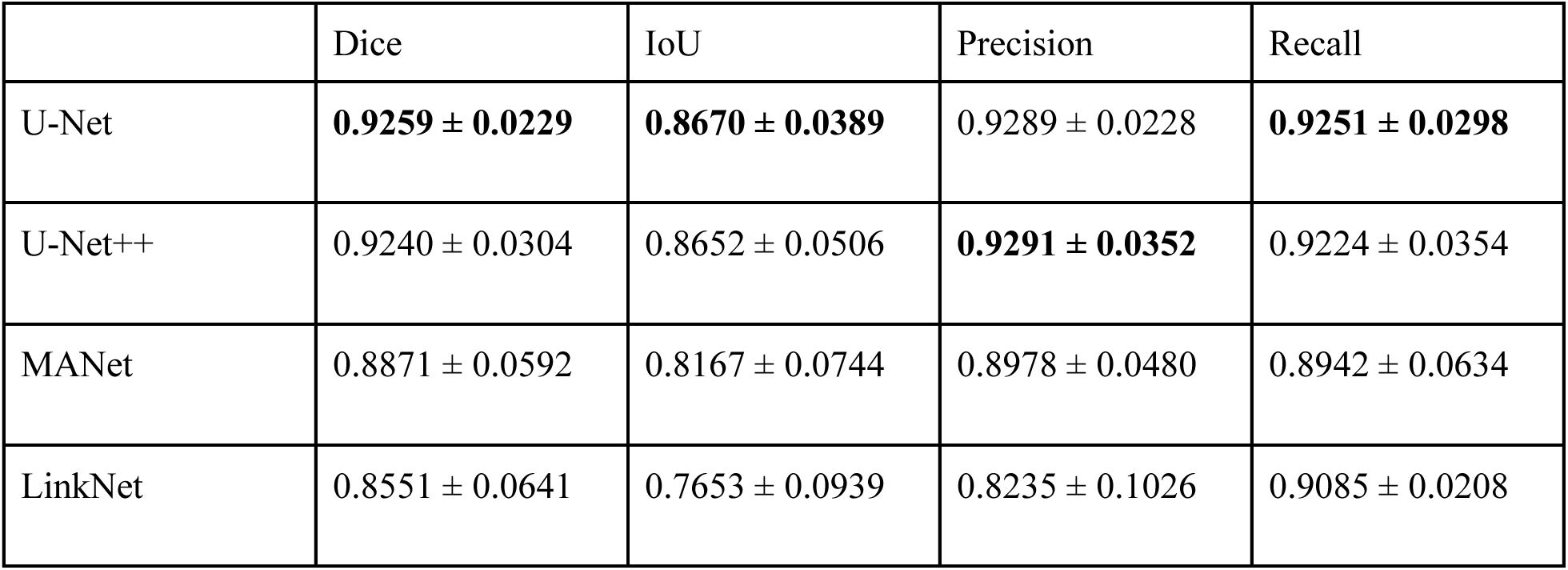
Model performance for GFAP Segmentation.

|  | Dice | IoU | Precision | Recall |
| --- | --- | --- | --- | --- |
| U-Net | **$0.9259 \pm 0.0229$** | **$0.8670 \pm 0.0389$** | $0.9289 \pm 0.0228$ | **$0.9251 \pm 0.0298$** |
| U-Net++ | $0.9240 \pm 0.0304$ | $0.8652 \pm 0.0506$ | **$0.9291 \pm 0.0352$** | $0.9224 \pm 0.0354$ |
| MANet | $0.8871 \pm 0.0592$ | $0.8167 \pm 0.0744$ | $0.8978 \pm 0.0480$ | $0.8942 \pm 0.0634$ |
| LinkNet | $0.8551 \pm 0.0641$ | $0.7653 \pm 0.0939$ | $0.8235 \pm 0.1026$ | $0.9085 \pm 0.0208$ |

### 3.2 IBA1 Segmentation

As shown in Table 4, all four models performed better on the IBA1 segmentation compared to the GFAP. U-Net++ achieved the highest Dice and IoU scores, along with leading precision and recall, indicating excellent detection of IBA1-positive regions. Low standard deviations across all four metrics suggest consistent performance, likely aided by the larger number of IBA1 images. U-Net performed competitively, leading in all categories after U-Net++. Results are shown in Fig. 4. Of the four models, MANet had lower performance and greater variability but still had strong numbers for all four metrics.

**Figure 4.**
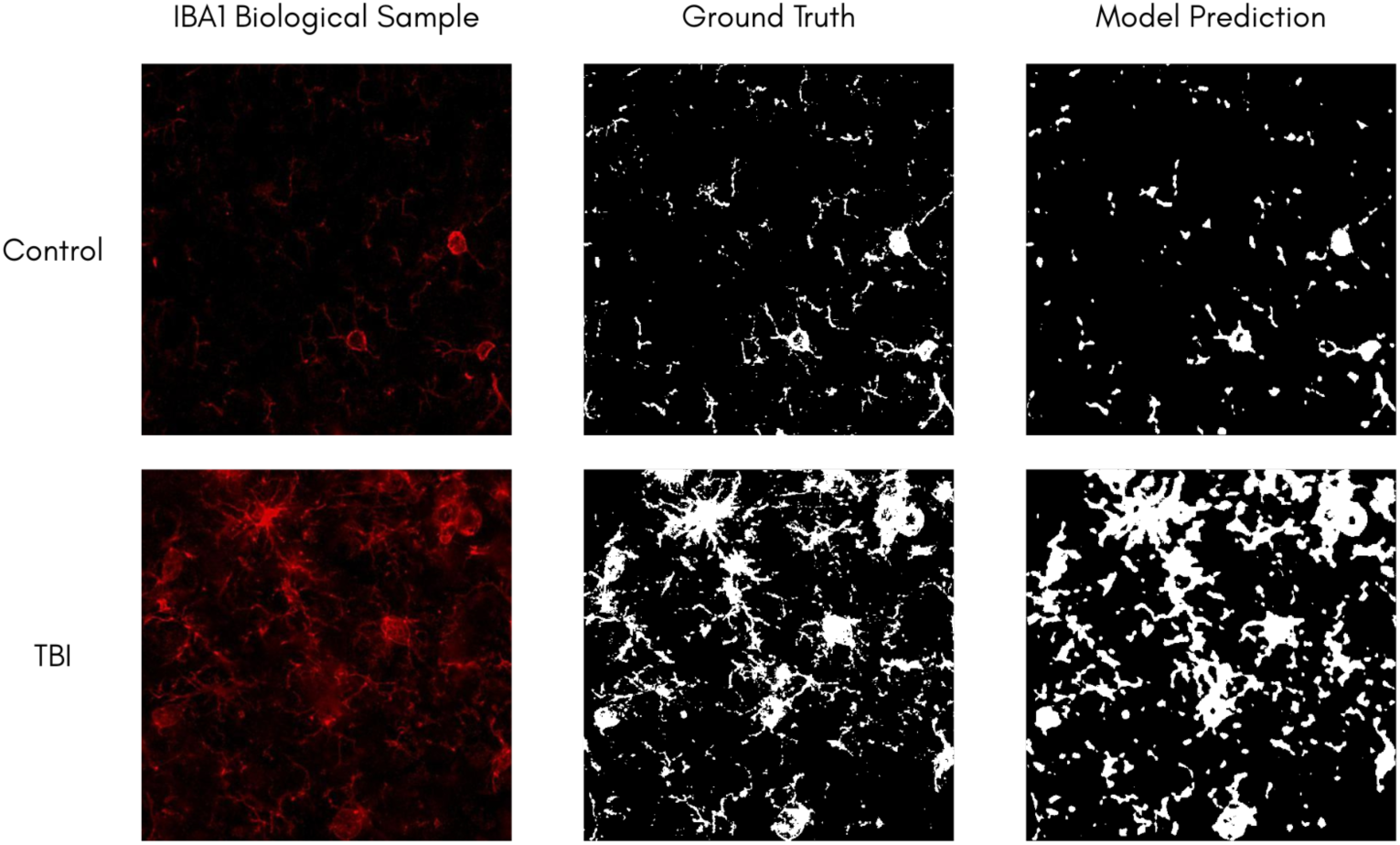
IBA1 segmentation prediction for TBI and Control, presented next to original tile and ground truth.

**Table 4.** Model performance for IBA1 Segmentation.

|  | Dice | IoU | Precision | Recall |
| --- | --- | --- | --- | --- |
| U-Net | $0.9603 \pm 0.0040$ | $0.9281 \pm 0.0049$ | $0.9661 \pm 0.0061$ | $0.9547 \pm 0.0036$ |
| U-Net++ | **$0.9676 \pm 0.0046$** | **$0.9418 \pm 0.0059$** | **$0.9721 \pm 0.0097$** | **$0.9635 \pm 0.0091$** |
| MANet | $0.9088 \pm 0.0417$ | $0.8427 \pm 0.0655$ | $0.9025 \pm 0.0501$ | $0.9221 \pm 0.0350$ |
| LinkNet | $0.9579 \pm 0.0042$ | $0.9234 \pm 0.0047$ | $0.9631 \pm 0.0090$ | $0.9531 \pm 0.0085$ |

### 3.3 QuPath GFAP Segmentation

Table 5 demonstrates that QuPath pixel analysis methods were moderately successful with segmenting immunofluorescent GFAP images for positive and negative regions. Artificial neural network (ANN) and logistic regression (LR) were significantly variable in trials with Dice, IoU, and Recall measurements, yet had stable performance with Precision measurements. Comparatively, the random forest (RF) algorithm had more stable results on other metrics. High precision with poor recall levels, especially for the logistic regression model, indicate that while positive predictions were largely correct, the model failed to detect the majority of true positive regions. All four U-Net algorithms in Section 3.1 performed significantly better across all metrics compared to traditional pixel classification (segmentation) methods within QuPath, with higher scores between 0.15 to 0.35. The only exception to this is the logistic regression’s precision, but its recall is significantly lower.

**Table 5.**
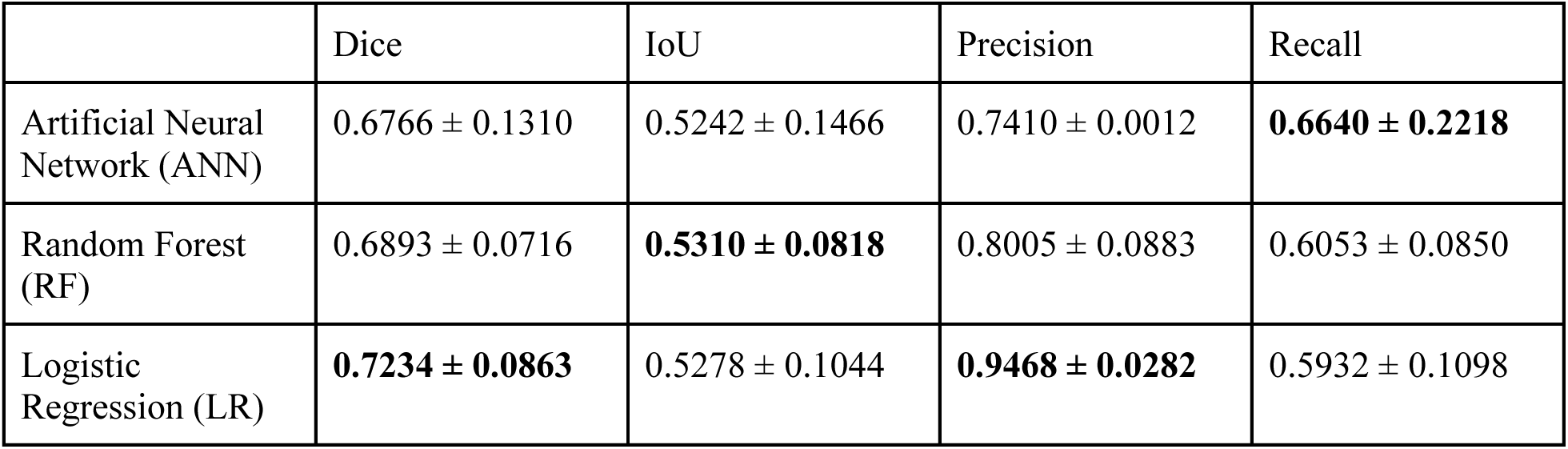
QuPath machine learning performance for GFAP Segmentation.

|  | Dice | IoU | Precision | Recall |
| --- | --- | --- | --- | --- |
| Artificial Neural Network (ANN) | $0.6766 \pm 0.1310$ | $0.5242 \pm 0.1466$ | $0.7410 \pm 0.0012$ | **$0.6640 \pm 0.2218$** |
| Random Forest (RF) | $0.6893 \pm 0.0716$ | **$0.5310 \pm 0.0818$** | $0.8005 \pm 0.0883$ | $0.6053 \pm 0.0850$ |
| Logistic Regression (LR) | **$0.7234 \pm 0.0863$** | $0.5278 \pm 0.1044$ | **$0.9468 \pm 0.0282$** | $0.5932 \pm 0.1098$ |

### 3.4 QuPath IBA1 Segmentation

Algorithms within QuPath mostly performed worse on the IBA1 dataset compared to the GFAP dataset, as seen in Table 6. While artificial neural network and logistic regression maintained scores from 0.53 to 0.76, which are acceptable scores, the random forest had notably lower scores on Dice, IoU, and Recall, while maintaining a high precision. Low variability indicates that this was a common trend across all ten folds, consistent with the pattern seen in the logistic regression’s performance in 3.3. U-Net machine learning models were still dominant compared to QuPath machine learning algorithms. Furthermore, it is important to note in addition to low performance compared to U-Net machine learning models, QuPath analysis was slower, with predictions taking several seconds compared to U-Nets completing predictions in milliseconds.

**Table 6.** QuPath machine learning performance for IBA1 Segmentation.

|  | Dice | IoU | Precision | Recall |
| --- | --- | --- | --- | --- |
| Artificial Neural Network | **0.6818 ± 0.0177** | **0.5311 ± 0.0217** | 0.7661 ± 0.0098 | 0.6234 ± 0.0313 |
| Random Forest | 0.4382 ± 0.0243 | 0.3187 ± 0.0276 | **0.9056 ± 0.0178** | 0.3266 ± 0.0211 |
| Logistic Regression | 0.6850 ± 0.0963 | 0.5282 ± 0.1113 | 0.7520 ± 0.1021 | **0.6367 ± 0.1083** |

### 3.5 GFAP Classification

Results in Table 7 show that across the three CNN classifiers, ResNet50 achieved the strongest overall performance, with the highest mean accuracy, sensitivity, and AUC-ROC, while maintaining competitive specificity and F1 Score. Swin_T outperformed other models in F1 Score, indicating balanced precision and recall. MaxViT, although strong, showed lower performance compared to the other models that do not rely on hybrid architectural design. Overall, both standalone architectures outperformed MaxViT, with ResNet50 providing the most consistent and accurate GFAP classification.

**Table 7.** Model performance for GFAP Classification.

|  | Accuracy | Sensitivity | Specificity | F1 Score | AUC-ROC |
| --- | --- | --- | --- | --- | --- |
| ResNet50 | **0.9456 ± 0.0537** | **0.9800 ± 0.0450** | 0.9092 ± 0.1065 | 0.9304 ± 0.0558 | **0.9933 ± 0.0129** |
| Swin_T | 0.9426 ± 0.0503 | 0.9467 ± 0.0612 | **0.9400 ± 0.0843** | **0.9357 ± 0.0528** | 0.9850 ± 0.0239 |
| MaxViT | 0.9283 ± 0.0470 | 0.9467 ± 0.0613 | 0.9147 ± 0.0817 | 0.9204 ± 0.0515 | 0.9860 ± 0.0192 |

### 3.6 IBA1 Classification

As shown in Table 8, IBA1 classification results were lower and more variable than GFAP. MaxViT showed the strongest performance across all five metrics, outperforming by 0.06 to 0.10 per metric. MaxViT also offered the most consistent performance, with lower variance in every single metric except for accuracy, where ResNet50 was less accurate but slightly more consistent. ResNet50 and Swin_T achieved similar results on all five metrics, with ResNet50 slightly leading in every single metric except AUC-ROC. In summary, IBA1 classification is more challenging than GFAP, but MaxViT’s advanced architecture contributes to the best-balanced performance.

**Table 8.** Model performance for IBA1 Classification.

|  | Accuracy | Sensitivity | Specificity | F1 Score | AUC-ROC |
| --- | --- | --- | --- | --- | --- |
| ResNet50 | 0.7366 ± 0.0414 | 0.7582 ± 0.1068 | 0.7065 ± 0.1266 | 0.7668 ± 0.0523 | 0.8278 ± 0.0396 |
| Swin_T | 0.7291 ± 0.0614 | 0.7492 ± 0.1514 | 0.7009 ± 0.1589 | 0.7565 ± 0.0769 | 0.8279 ± 0.0563 |
| MaxViT | **0.8118 ± 0.0425** | **0.8170 ± 0.0707** | **0.8045 ± 0.0496** | **0.8334 ± 0.0434** | **0.9008 ± 0.0348** |

### 3.7 Paired IBA1 & GFAP Classification

The integration of IBA1 and GFAP into a multimodal input enhanced classification performance compared to single-marker models, as seen in Table 9. All three architectures benefited from the integration of astrocytic and microglial information, which provided richer spatial and contextual cues for distinguishing TBI from control samples. Scores are significantly better compared to IBA1, and Swin_T and MaxViT trained on paired samples beat GFAP on every single metric. ResNet50 trained on paired samples shows slightly lower performance in sensitivity and AUC-ROC, but has a better F1-Score, indicating better balance of precision and recall.

**Table 9.** Model performance for paired GFAP-IBA1 Classification.

|  | Accuracy | Sensitivity | Specificity | F1 Score | AUC-ROC |
| --- | --- | --- | --- | --- | --- |
| ResNet50 | 0.9480 ± 0.0433 | 0.9600 ± 0.0699 | 0.9378 ± 0.0540 | **0.9499 ± 0.0437** | 0.9869 ± 0.0152 |
| Swin_T | **0.9489 ± 0.0968** | 0.9500 ± 0.1333 | **0.9478 ± 0.0768** | 0.9466 ± 0.1034 | **0.9909 ± 0.0252** |
| MaxViT | 0.9431 ± 0.0570 | **0.9689 ± 0.0502** | 0.9156 ± 0.1116 | 0.9462 ± 0.0519 | 0.9903 ± 0.0211 |

## 4. Discussion

In this study, we performed a comprehensive evaluation of multiple deep learning architectures for the segmentation and classification of GFAP and IBA1 immunofluorescence images, considering both single-marker and dual-marker conditions. Across all experiments, results consistently demonstrate that modern deep learning models can achieve high performance, especially when leveraging dual-marker inputs.

### 4.1 Segmentation Performance Across GFAP and IBA1

Segmentation results indicate strong performance among all architectures for both GFAP and IBA1, with the best Dice scores exceeding 0.92 for GFAP and 0.96 for IBA1. U-Net consistently achieved the highest accuracy in GFAP, outperforming the other three models in Dice, IoU, and Recall metrics. The more advanced U-Net++ achieved the strongest overall performance in IBA1, leading in performance across all four metrics, with these findings confirming the robustness of such models for biomedical segmentation tasks. The results also suggest that more complex architectures may not always provide significant advantage for certain imaging tasks. This was seen in the performance of U-Net, which matched or exceeded U-Net++ across most GFAP metrics with the exception of precision. IBA1 segmentation exhibited slightly higher overall performance compared to GFAP, benefiting from a larger training dataset despite more complex morphological structures. LinkNet demonstrated the lowest performance for GFAP. MANet demonstrated the lowest performance for IBA1, with higher standard deviations suggesting reduced consistency, which demonstrates a potential trade-off with the sensitivity of attention mechanisms.

### 4.2 Classification Performance Across GFAP and IBA1

GFAP single-channel classification produced strong results across all models, with ResNet50 achieving the best overall performance. High sensitivity and AUC-ROC values indicate effective separation between positive and negative classes, even with a relatively small dataset. In contrast, IBA1 single-channel classification was significantly more challenging, with no model averaging higher than 0.9008 on any metric. On this dataset, MaxViT demonstrated significantly higher scores with greater consistency on every metric. This may reflect the subtle staining patterns of IBA1, which complicate discrimination between negative and weakly positive samples. This also suggests that MaxViT’s hybrid convolutional-transformer architecture better captures contextual cues inherent in IBA1 morphology.

### 4.3 Benefits of Dual-Marker Inputs

Introducing dual-channel inputs combining IBA1 and GFAP significantly improved classification performance across all models. Accuracy, F1 Score, and AUC-ROC values increased markedly, with Swin_T achieving the strongest accuracy, although ResNet50 achieved the strongest F1 Score. Improvements in certain metrics in the two-channel trial were particularly notable; for example, ResNet50 sensitivity increased from 0.7582 (IBA1 alone) to 0.9600 (IBA1 and GFAP). These findings underscore the value of integrating complementary biological information, as astrocytic cues help reduce false positives in microglial classification. Overall, dual-marker integration enhances model robustness and reliability in medical imaging analysis.

## 5. Conclusion

This work presents a deep learning framework that demonstrates the benefits of incorporating multiple biomarkers for improved machine learning analysis of histological images. Our findings show that deep learning models can achieve high performance in both segmentation and classification of GFAP and IBA1 immunofluorescence images. Simpler models, like U-Net and ResNet50, performed competitively with more advanced architectures like U-Net++, MaxViT, and Swin_T. Differences between GFAP and IBA1 performance highlight the influence of staining characteristics on model behavior, with IBA1 posing a greater challenge due to subtle morphological variability.

Combining GFAP and IBA1 in a dual-channel input significantly improved classification accuracy, emphasizing the importance of multimodal approaches in neuroinflammation and glial activation studies. Although demonstrated here in the context of TBI, this framework can be extended beyond TBI and applied to other neurological conditions. Future work may explore additional biomarkers, expand to diverse injury models, and evaluate performance across imaging modalities beyond immunofluorescence. These findings support the integration of multi-channel strategies for disease modeling, drug testing, and neurological assessment.

